# Effect of seizures on the severity of myelin vacuolization in a mouse model of megalencephalic leukoencephalopathy with subcortical cysts

**DOI:** 10.1101/2024.04.23.590685

**Authors:** Eelke Brouwers, Marjolein Breur, Maria Susana Jorge, Huibert D. Mansvelder, Marjo S. van der Knaap, Marianna Bugiani, Rogier Min

**Author notes:** These authors jointly supervised this study (Marianna Bugiani, Rogier Min).

## Abstract

**Objective:** Megalencephalic leukoencephalopathy with subcortical cysts (MLC) is a white matter disease characterized by myelin vacuolization and swollen perivascular astrocyte processes. Neuronal activity has been implicated as primary cause of myelin vacuolization and astrocyte swelling. Since acute and excessive increases in neuronal activity occur in MLC during seizures, we investigated whether seizure activity leads to the acute development of myelin vacuoles and swollen astrocyte processes.

**Methods:** *Glialcam*-null mice (an established MLC mouse model) and wild-type mice received repeated i.p. low dose (5 mg / kg) kainic acid (KA) injections until severe seizures developed. Following a 60-minute period of severe seizure activity, mice were terminated and brains were fixed and processed. Brain tissue was analyzed for myelin vacuolization and astrocyte process thickness using H&E and GFAP stains, respectively.

**Results:** Repeated low-dose injections of KA resulted in prolonged severe seizure activity in mice of both genotypes. Total amount of seizure activity was comparable between *Glialcam*-null and wild-type mice. KA-induced severe seizure activity did not significantly increase myelin vacuolization in either *Glialcam*-null or wild-type mice. The width of perivascular astrocyte processes was also not affected by severe seizure activity.

**Interpretation:** We show that (i) repeatedly injecting a low dose of KA provides the opportunity to regulate seizure development and generate severe seizures in both wild-type and seizure-sensitive *Glialcam*-null mice, and that (ii) the two major pathological features of MLC, myelin vacuolization and swollen astrocyte endfeet, are not acutely aggravated in response to KA-induced severe seizure activity.

## INTRODUCTION

Many axons in the brain are ensheathed by multiple layers of myelin, produced by oligodendrocytes. Myelination greatly improves the speed and efficiency of electrical signal conduction between neurons by enabling saltatory conduction between adjacent nodes of Ranvier (Hartline & Colman, 2007; Tasaki, 1939). In addition, myelinating oligodendrocytes provide the axons with metabolic support and play a critical role in buffering expelled potassium (Funfschilling et al., 2012; Larson et al., 2018). The structure of myelin affects the speed of action potential conduction, with myelin thickness and the distribution and length of nodes and internodes precisely regulated (Arancibia-Carcamo et al., 2017; Waxman, 1980). Additionally, the nanoscale properties of the periaxonal space (the space between axon and innermost myelin layer) crucially contributes to the speed of saltatory conduction (Cohen et al., 2020). The importance of myelin structure for healthy brain functioning is evident from the wide variety of neurological diseases associated with myelin changes.

Diseases associated with myelin dysfunction can be broadly placed into three categories: demyelinating diseases, hypomyelinating diseases and diseases with myelin vacuolization. Demyelinating diseases are characterized by the breakdown of myelin, as occurs in multiple sclerosis (MS) and several other acquired neurological diseases (Lassmann, 2001). Demyelination may also be caused by genetic defects, such as in metachromatic leukodystrophy (van der Knaap & Bugiani, 2017). In hypomyelinating diseases the production of myelin is transiently or permanently reduced (for review see (Barkovich & Deon, 2016; Pouwels et al., 2014). Finally, in myelin vacuolating diseases, the myelin is present, but contains vacuoles. Such vacuolization can sometimes, but certainly not always, result in demyelination (Duncan & Radcliff, 2016; van der Knaap & Bugiani, 2017). Importantly, the pathological mechanisms and dynamics underlying myelin vacuolization are not completely understood.

Several studies have shown a relation between the development of myelin vacuoles and neuronal activity (Blanz et al., 2007; Menichella et al., 2006; Wurtz & Ellisman, 1986). For example, high frequency stimulation of healthy frog dorsal root fibers results in the acute development of severe myelin vacuolization (Wurtz & Ellisman, 1986). This illustrates that myelin vacuoles can develop when healthy axons are overloaded with neuronal activity. Interestingly most myelin vacuoles disappear within 30 minutes after the stimulus, indicating that myelin vacuolization can be reversible. Observations in transgenic mouse models for neurological diseases characterized by myelin vacuolization also support activity dependence. For example, in the *Clcn2*-null mouse, which shows extensive myelin vacuolization, the optic nerve is spared from such vacuolization (Blanz et al., 2007). This mouse is blind due to extensive retinal degeneration, meaning that the optic nerve is electrically silent. This supports the idea that action potential firing is a prerequisite for the development of myelin vacuolization. Similarly, in double knockout mice for the gap junction subunits Cx32/Cx47, myelin vacuolization of the optic nerve develops upon eye opening, when the optic nerve becomes electrically active. Blocking optic nerve activity with TTX or by covering the eye prevents the development of myelin vacuoles (Menichella et al., 2006). What remains unclear is the timescale of activity-dependent myelin vacuolization in the brain. While acute development and quick resolution of vacuoles has been reported in the peripheral nervous system (PNS) of the frog, activity-dependent vacuolization in the optic nerve of dKO Cx32/Cx47 was observed to occur on a time-scale of days.

The mechanism underlying activity dependent myelin vacuolization is likely linked to action potential mediated shifts in ions, and accompanying alterations in osmotic pressure. Neuronal activity induced action potential propagation along myelinated axons causes local increases of potassium in the periaxonal space underneath the myelin (David, Barrett, & Barrett, 1993). Periaxonal potassium build-up has disastrous consequences for the maintenance of action potential firing and the excitability of the neuron, and in extreme situations can lead to conduction failure or onset of seizures (Brazhe, Maksimov, Mosekilde, & Sosnovtseva, 2011). It is therefore crucial that periaxonal concentrations of potassium are kept low. The mechanisms of periaxonal potassium clearance are not completely understood. It has been suggested that potassium is taken up by the innermost myelin layers and is further distribution away from the axon, into the network of coupled astrocytes. This model has been extensively described by Rash, and emphasizes the role of gap junctional coupling between successive myelin layers and between oligodendrocytes and astrocytes (i.e. the panglial syncytium) in long-distance distribution of potassium (Rash, 2010). Oligodendrocyte and astrocyte (O/A) gap junctions are formed by connexin Cx47/Cx43 and Cx32/Cx30 pairs, whereas connexin Cx32 alone connects adjacent myelin layers (Kamasawa et al., 2005; Orthmann-Murphy, Freidin, Fischer, Scherer, & Abrams, 2007). Decreased drainage possibilities due to loss of certain proteins, such as the connexins Cx47 and Cx32, would then lead to an accumulation of periaxonal or intramyelinic potassium, especially during excessive neuronal firing. Osmotic swelling due to the accompanying osmotic disturbance could potentially result in myelin vacuolization.

Restrictions in potassium buffering from the axon towards the astrocyte network are not limited to loss of function of oligodendrocyte / myelin proteins. Dysfunction of astrocytes, which form an important part of the panglial syncytium, can likewise hamper the distribution of potassium and thereby affect myelin integrity. This is clearly illustrated in the rare heritable white matter disease Megalencephalic Leukoencephalopathy with subcortical Cysts (MLC), where loss of the astrocyte specific protein MLC1 not only leads to the swelling of perivascular astrocyte processes but also to severe myelin vacuolization. Myelin vacuolization in MLC is functionally unrelated to myelin generation and oligodendrocyte health. Patients as well as mouse models for MLC show normal myelin development, with no difference in early myelin appearance and adult myelin levels. Progressive vacuolization occurs as soon as myelin is laid down, but is not primarily accompanied by death of oligodendrocytes, axons or neurons (Dubey et al., 2015; van der Knaap, Barth, Vrensen, & Valk, 1996). Acute increases in neuronal activity occur in MLC patients due to the presence of epileptic seizures (Dubey et al., 2018). MLC mice are more prone to seizure activity, but do not show overt seizures. It is unclear what the effect of seizures is on the development of myelin vacuolization in MLC. We hypothesize that severe seizure activity acutely contributes to vacuole formation due to ionic and osmotic disturbance, and that this is readily observed in the MLC brain because of impaired functioning of the panglial syncytium. To test this hypothesis, we induced severe seizure activity in both wildtype and MLC mice by administration of the neuroexcitatory compound kainic acid (KA). We studied the amount of myelin vacuolization immediately following a period of severe seizure activity as well as after a short recovery period.

## MATERIALS & METHODS

### Mice

All experiments were approved by the Institutional Animal Care and Use Committee of the VU University, Amsterdam, and were in strict compliance with animal welfare policies of the Dutch government. *Glialcam*-null mice, of which the generation and characterization was described previously, were used as MLC model (Bugiani, Dubey et al. 2017). For this study, a total of 12 male wild-type and 12 male *Glialcam*-null mice aged 3-5 months were included. The genotype was defined by PCR and confirmed by re-genotyping after recording. Mice were housed in a temperature- and humidity-controlled room on a reversed 12h light-dark cycle (lights off at 9 am) with *ad libitum* access to food and water.

### Repeated low dose kainic acid injections

For seizure recording sessions, mice were transferred to a Perspex experimental cage (containing bedding from their home cage), allowing optimal behavioral observation. Video monitoring was performed throughout the experiment (software: iSpy v5.8.6.0). Mice were allowed familiarization with the new environment and assessed for ‘baseline’ behavior for 10 minutes. To induce controlled epileptic seizures, we used a repeated low dose kainic acid injection regime (Tse, Puttachary, Beamer, Sills, & Thippeswamy, 2014). Mice received multiple intraperitoneal (i.p.) injections of either saline or kainic acid (KA) in doses of 5 mg/kg. KA (1mg/ml) was dissolved in sterile saline and prepared fresh on the day of the experiment. The first injection was given following the baseline period. Subsequent injections followed at 30-minute intervals. Mice in the KA group continued to receive injections until a first seizure classified as stage 4 or higher occurred (see ‘behavioral assessment’ below for details on seizure classification). After this first seizure classified as stage 4 or higher, mice were video monitored for an additional 60 minutes. Mice in the saline group received a total of 4 injections (5ml/kg; equal volume as in the KA group). After the fourth injection, these mice were also video monitored for an additional 60 minutes. At the end of the final 60-minute period all mice were decapitated under isoflurane anesthesia and brains were extracted. Brains were immersion fixed in 4% PFA for further histological processing. In a subset of mice, morphological alterations after full recovery from seizures were also assessed. The KA injection regime was similar to that described above. However, at the end of the final 60-minute period these mice received a 300 mg/kg valproic acid (VPA) injection i.p. to reduce seizure activity. VPA was prepared as a 30mg/ml solution and dissolved in sterile saline. VPA injection was followed by a 6-hour recovery period. At the end of this period mice were sacrificed and brains were extracted as described above.

### Behavioral assessment

Seizure severity was scored using the Racine scale described by Tse and colleagues (Tse et al., 2014). For clarification:

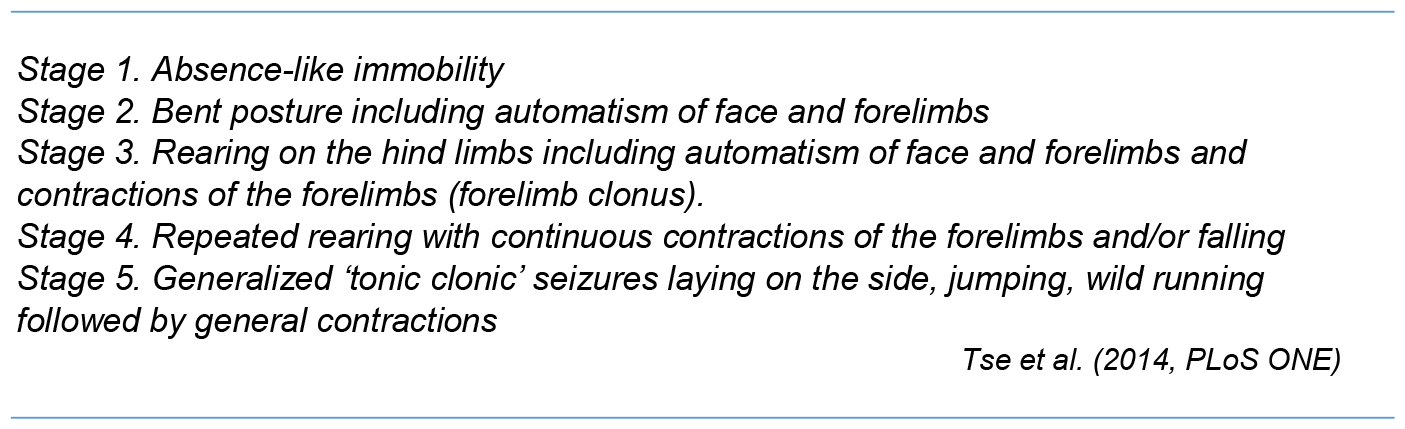

Offline video analyses and scoring of behavior was performed by a trained observer blind for genotype. One wild-type mouse and one *Glialcam*-null mouse experienced a severe seizure which was missed during online assessment, and therefore received one KA injection too much. We used the offline assessment for the analysis of the number of injections required to develop severe seizures. For most mice, maximum seizure score per 5-minute bins was reported. A subgroup of mice was analyzed in more detail, where the exact time point of every transition in seizure severity was defined (**Fig. 1**).

**FIGURE 1:**
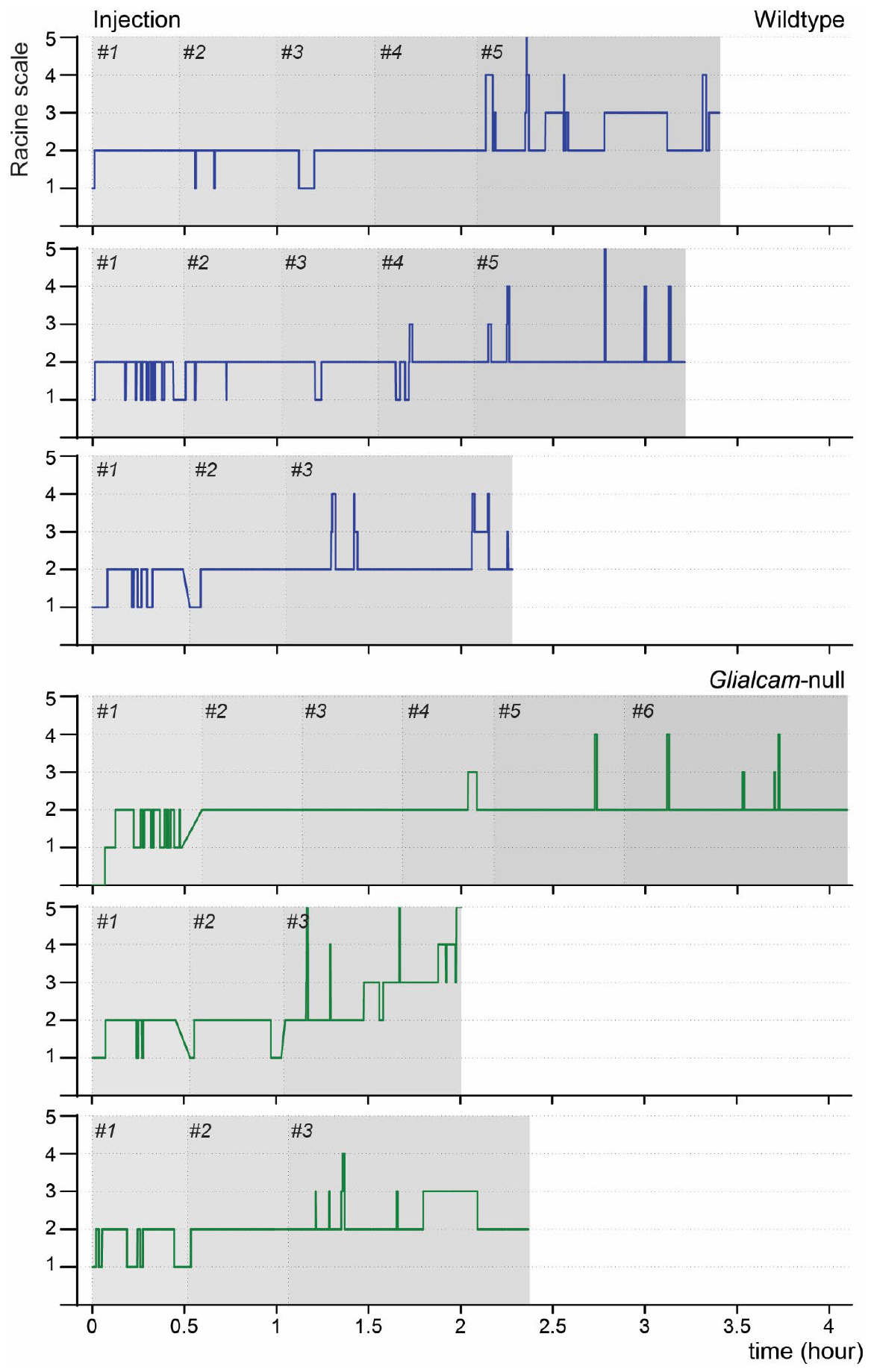
Seizure development over time in response to repeated low-dose KA injections. The graph presents example traces of three wild-type mice (top, blue) and three *Glialcam*-null mice (bottom, green). Number (#) of injections received and corresponding time points are indicated on the top of each graph and visualized by shaded areas.

### Quantification of astrocyte process thickness and myelin vacuoles

Formalin fixed paraffin embedded brain tissues were sliced into 5um thick sections and stained against glial fibrillary acidic protein (GFAP) or with H&E to visualize perivascular astrocytic cell processes and myelin respectively. Quantification of astrocyte morphology and vacuoles was performed blind to the genotype and treatment using ImageJ. The maximum width of astrocyte processes was measured using 40x magnified images. The percentage of cross-sectioned area occupied by vacuoles was quantified in 4 images of the cerebellar folia.

### Statistical analysis

Behavioral data were analyzed with the unpaired t-test (cumulative severity score and time to onset of first severe seizure) and the Mann-Whitney test (number of injections). For morphological data analysis we used a nested one-way ANOVA with six preselected multiple comparisons, Sidak corrected:

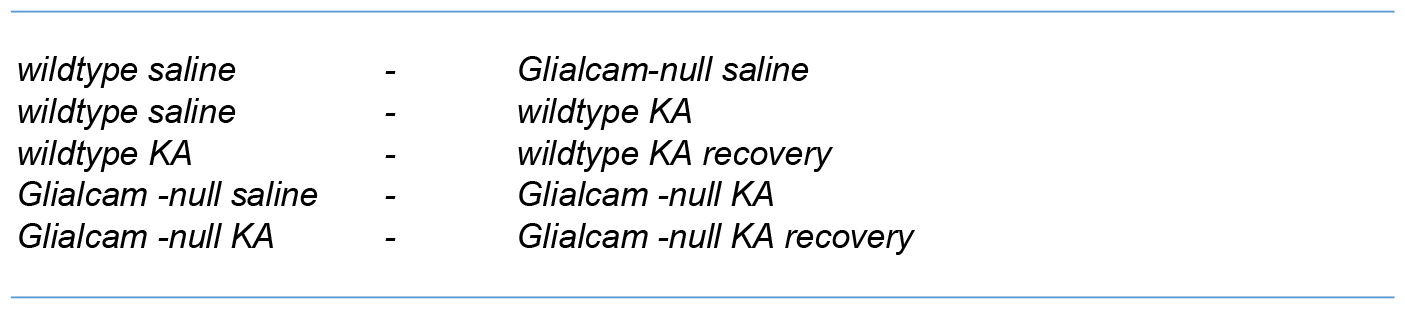

Astrocyte data were log transformed to meet the normality distribution condition. Data were processed with Prism v8.2 (GraphPad software). Probability values <0.05 were considered significant. Data are presented as mean ± SEM.

## RESULTS

### Equalizing seizure severity among wild-type and Glialcam-null mice by repeatedly injecting a low dose of kainic acid

MLC mice have a lower threshold for developing KA-induced seizures compared to wild-types (Dubey et al., 2018). With a single dose of 10 mg / kg KA, MLC mice developed severe seizures (Racine score 4-5), while wild-type mice showed mainly mild seizures (Racine score 2-3). Pilot experiments, in which we looked at brain tissue following seizures, suggested that the pathological features of MLC (myelin vacuolization in cerebellar white matter and thickness of astrocyte endfeet) increase in both wild-type and MLC mice after induced seizures (data not shown). However, from these experiments it was unclear whether the observed elevation differs between MLC and wild-type mice, since MLC mice experienced more severe seizures. This precluded drawing conclusions about seizure-induced changes in astrocyte and myelin morphology between MLC and wild-type mice.

In the present study, we opted for an approach in which both *Glialcam*-null and wild-type mice eventually experienced severe seizures for a similar duration. This was achieved by adjusting the amount of administered KA based on the course of seizure activity. Mice repeatedly received a low dose of KA (5 mg/kg) at 30-minute intervals, until a first severe seizure (Racine score 4 or 5) was observed. From then on, mice were monitored for an additional 60 minutes without further injections, after which they were sacrificed and the brains were extracted for histological analysis. The effectiveness of this repeated low-dose method for controlling seizures in wild-type animals has been described previously (Tse et al., 2014; Umpierre et al., 2016). We show here that it was also successful for equalizing seizure severity among MLC and wild-type animals. An example of the course of the experiment and the development of seizures for 3 wild-type mice and 3 *Glialcam*-null mice is shown in **figure 1**. In this example, severe seizures were observed from the third injection onwards.

Overall, *Glialcam*-null animals did not differ from wild-type mice in the amount of KA injections required to induce severe seizures (wild-type: 4.11 ± 0.26 injections, n = 9; *Glialcam*-null: 3.67 ± 0.29 injections, n = 9; *P* = 0.29; **Fig. 2b**). Similarly, no difference was found in the time between the first KA injection and the first severe seizure (wild-type: 106.1 ± 7.93 minutes, n = 9; *Glialcam*-null: 98.33 ± 12.67 minutes, n = 9; *P* = 0.61; **Fig. 2c**). To compare the development of seizures over time, we defined the maximum seizure stage per 5-minute time bin for each mouse. Average scores per genotype are shown in **figure 2a**. Each star represents the time point of an individual mouse reaching a first Racine stage 4/5. We subsequently summed the values from the 5-min bin analysis per mouse to calculate the cumulative seizure severity (**Fig. 2d, left**). From this we could conclude that the total amount of experienced seizure activity did not differ between groups (cumulative seizure score: wild-type: 79.89 ± 3.91, n = 9; *Glialcam*-null: 78.44 ± 6.82, n = 9; *P* = 0.86). We also examined whether MLC mice experience more severe seizures from the moment they enter stage 4/5 by calculating the cumulative score of the last 60 minutes following the final KA injection. Again, no clear difference was observed in seizure severity (wild-type: 35.44 ± 2.82, n = 9; *Glialcam*-null: 41.00 ± 3.76, n = 9; *P*= 0.25). We conclude that the repeated low KA dose injection regime is an appropriate method for equalizing the amount of experienced seizure activity between *Glialcam*-null and wild-type mice.

**FIGURE 2:**
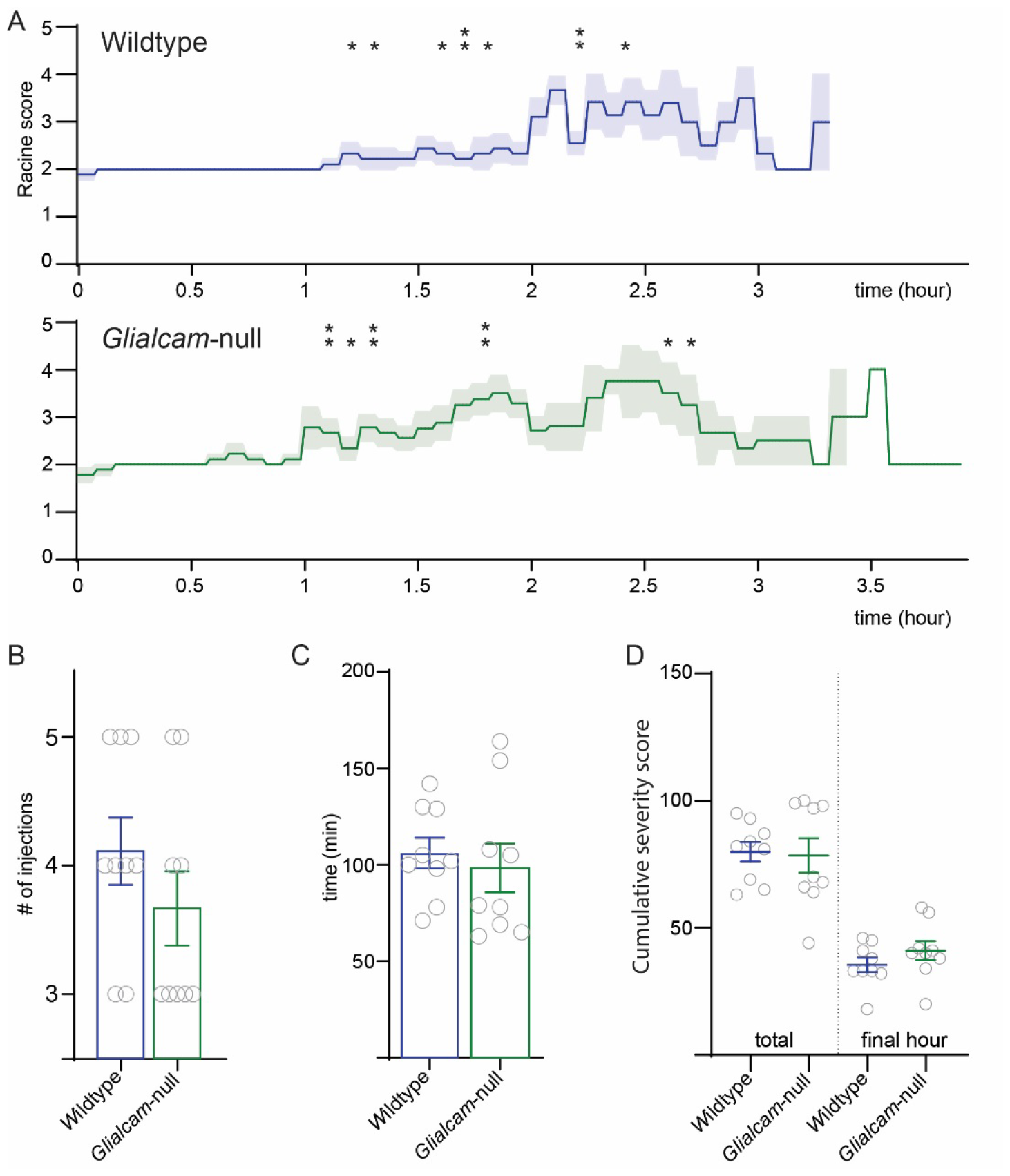
Comparison of cumulative seizure severity and number of received KA injections between wild-type and *Glialcam*-null mice. (A) Averaged Racine scores per 5 minutes for 9 wild-type (top, blue) and 9 *Glialcam*-null (bottom, green) mice. The time point of the first severe seizure (Racine stage 4/5) for individual mice is indicated by an asterisk. Shaded areas represent the SEM. (B) Number of injections needed to develop severe seizures and (C) time to first severe seizure did not differ between wild-type (blue) and *Glialcam*-null (green) mice. (D) Cumulative seizure severity during 60 minutes and over the total duration of the experiment were also not different between wild-type (blue) and *Glialcam*-null (green) mice. Open circles show individual mice. Bars indicate mean ± SEM.

### Kainic acid-induced seizures do not lead to acute myelin vacuolization in cerebellar white matter

Similar to patients, MLC mouse models are characterized by myelin vacuolization in the white matter (Bugiani M, Dubey M, et al. 2017). While patients show prominent vacuolization of the cerebral white matter, vacuolization in mice, which have virtually no cerebral hemispheric white matter, is most prominent in the cerebellar white matter (Dubey et al., 2015). Since myelin vacuolization has been suggested to depend on neuronal activity, we investigated whether seizures lead to acute development of cerebellar myelin vacuoles. In line with previous studies, we observed minimal presence of myelin vacuoles in saline injected wild-type mice, while saline injected *Glialcam*-null mice showed an increased vacuolated total area (percentage vacuolated area: wild-type: 0.87 ± 0.27 %, n = 12, *N* = 3; *Glialcam*-null: 6.37 ± 1.89 %, n = 12, *N* = 3; *P* = 0.0374, one-way ANOVA; **Fig. 3**). Next, we measured vacuolization in mice that received repeated low-dose KA injections, where mice were sacrificed and brains fixed 1 hour after the final KA injection. Repeated low-dose KA-induced seizures did not acutely affect the degree of vacuolization in wild-type mice or *Glialcam*-null mice (wild-type saline: 0.87 ± 0.27 %, n = 12, *N* = 3 vs wild-type KA: 1.69 ± 0.44 %, n = 16, *N* = 4; *P* = 0.99; *Glialcam*-null saline: 6.37 ± 1.89 %, n = 12, *N* = 3 vs *Glialcam*-null KA: 6.9 ± 0.86 %, n = 20, *N* = 5; *P* = 0.99, one-way ANOVA; **Fig. 3**). In a separate experimental group, mice received the same repeated low-dose KA injections followed by a 1 hour waiting period. After this, seizures were halted by injection of valproic acid (VPA) followed by a subsequent seizure free period of 6 hours. After this recovery period mice were sacrificed and brains were fixed. In this recovery group we observed that vacuolization had slightly increased in *Glialcam*-null mice, while such an increase was not observed in wild-type mice (wild-type KA: 1.69 ± 0.44 %, n = 16, *N* = 4 vs wild-type KA recovery: 2.97 ± 0.70 %, n = 16, *N* = 4; *P* = 0.96; *Glialcam*-null KA: 6.91 ± 0.86 %, n = 20, *N* = 5 vs *Glialcam*-null KA recovery: 11.26 ± 2.2 %, n = 16, *N* = 4; *P* = 0.049, one-way ANOVA; **Fig. 3**). Based on these results, we conclude that seizure activity did not lead to an acute increase in vacuolization of the cerebellar white matter. However, a small increase in white matter vacuolization after the seizures stopped was only observed in the *Glialcam*-null recovery group, suggesting slow development of myelin vacuoles following the initial seizure period in these mice.

**FIGURE 3:**
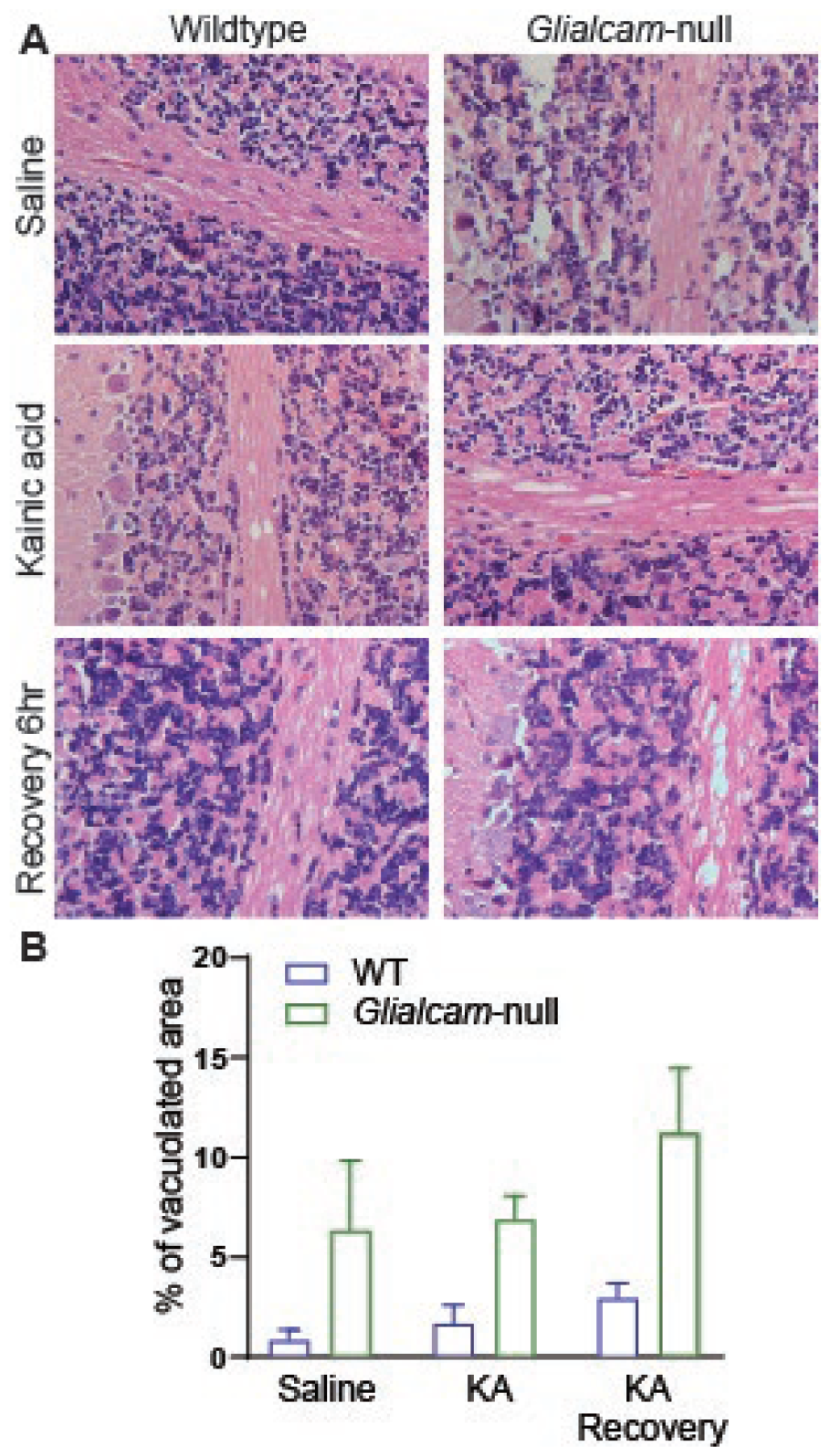
Myelin vacuolization in the cerebellar white matter of wild-type and *Glialcam*-null mice following KA induced severe seizure activity. (A) Representative example images of hematoxylin and eosin staining of the cerebellar white matter of wild-type (left) and *Glialcam*-null (right) mice upon saline injections (top), KA injections (middle) and KA injections followed by a VA injection and 6-hour recovery (bottom). (B) KA induced seizures had no effect on the percentage of vacuolated area in both wild-type (blue) and *Glialcam*-null (green) mice. Bars indicate mean ±.

The above results were statistically analyzed using one-way ANOVA with multiple comparison correction, in line with previous studies analyzing similar data (Bugiani et al., 2017; Dubey et al., 2015). We analyzed amount of vacuolization in four histological samples taken from each mouse, and treated the percentage vacuolization in each sample as an independent observation. The use of a regular one-way ANOVA on this data entails a chance of type I error (Aarts, Verhage, Veenvliet, Dolan, & van der Sluis, 2014). Therefore, it has been suggested that a more appropriate way to analyze such data is by taking a nested statistical approach, where potential dependency between samples obtained from the same mouse is considered (Aarts et al., 2014). Analyzing the same data with a nested one-way ANOVA did not affect vacuolization measures, but led to the absence of significant differences in vacuolization between any of the groups. This indicated that our present study was underpowered for detecting subtle differences between groups. Therefore, although subtle differences in the amount of vacuolization could not be assessed in our data set because of the small sample size, we conclude that KA-induced seizure activity does not have a profound effect on cerebellar white matter vacuolization in either wild-type or *Glialcam*-null mice.

### Kainic acid-induced seizures do not cause morphological changes in astrocyte endfeet thickness

Perivascular astrocytes from MLC patients and MLC mice were previously found to have thicker cell processes abutting blood vessels (Bugiani et al., 2017; Dubey et al., 2015). Astrocyte processes in contact with blood vessels play an important role in the exchange of ions and water between the internal environment of the brain and the blood circulation. The decreased ability of MLC astrocytes to deal with water shifts and their impaired volume regulation is thought to cause the observed thickening of their processes. We investigated to what extent seizure activity influences the thickness of astrocyte processes abutting blood vessels in *Glialcam*-null and wild-type mice, by injecting mice with KA or saline. Unexpectedly, saline injected *Glialcam*-null mice did not show increased astrocyte process thickness compared to wild-type mice (wild-type: 1.83 ± 0.03 µm, n = 474, *N =* 3; *Glialcam*-null: 1.57 ± 0.02 µm, n = 575, *N* = 3; *P* = 0.99, nested One-way ANOVA; **Fig. 4**). This is in contrast with a previous study on untreated *Glialcam*-null mice. Our value for the process thickness of 3-month-old saline-injected *Glialcam*-null mice (1.57 ± 0.02 µm) corresponds to what was reported in this earlier study (1.64 ± 0.04 µm (Bugiani et al., 2017)). However, processes of 3-to 5-month-old saline injected wild-type mice in our study appeared much thicker than those of 12-month-old wild-type animals reported by Bugiani *et al* (Bugiani et al., 2017). These results suggest a previously unnoticed age-dependent alteration of astrocyte process thickness in wild-type mice.

**FIGURE 4:**
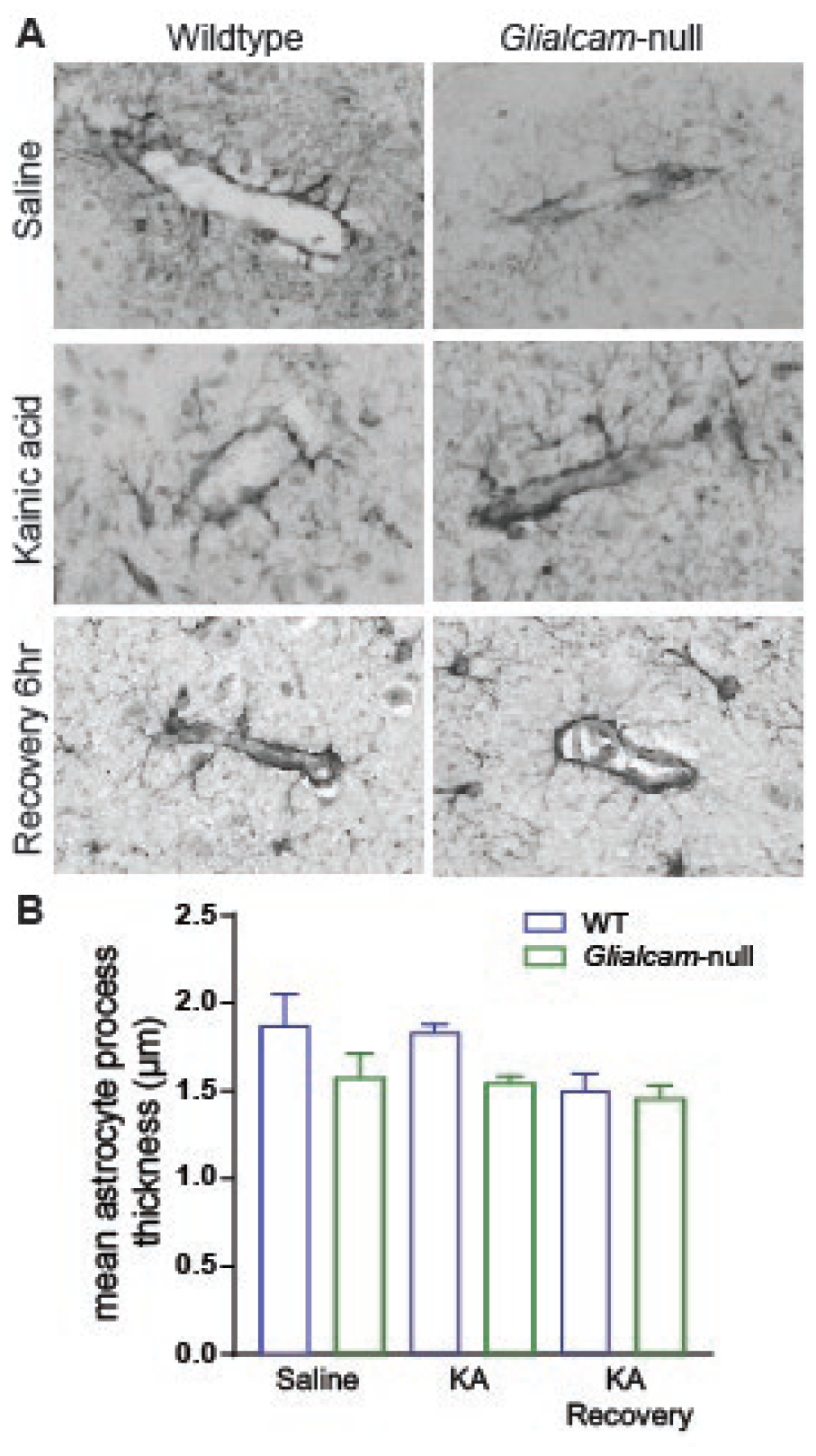
Astrocyte process thickness of wild-type and *Glialcam*-null mice following KA induced severe seizure activity. (A) Representative example images of Glial fibrillary acidic protein (GFAP) staining showing perivascular astrocytes abutting a blood vessel in wild-type (left) and *Glialcam*-null (right) mice upon saline injections (top), KA injections (middle) and KA injections followed by a VA injection and 6-hour recovery (bottom). (B) KA induced seizures had no effect on the thickness of perivascular astrocyte end feet in both wild-type (blue) and *Glialcam*-null (green) mice. Bars indicate mean ± SEM. KA = kainic acid.

Astrocyte process thickness was independent of seizure activity, since neither *Glialcam*-null nor wild-type mice showed changes in thickness after KA-induced seizures (wild-type saline: 1.83 ± 0.03 µm, n = 474, *N* = 3 vs wild-type KA: 1.83 ± 0.02 µm, n = 622, *N* = 4; *P* = >0.99; *Glialcam*-null saline: 1.57 ± 0.02 µm, n = 575, *N* = 3 vs *Glialcam*-null KA: 1.55 ± 0.02 µm, n = 1002, *N* = 5; *P* = >0.99, nested one-way ANOVA; **Fig. 4**). Additionally, no alterations of process thickness were observed in the recovery group (wild-type KA: 1.83 ± 0.02 µm, n = 622, *N* = 4 vs wild-type KA recovery: 1.51 ± 0.02 µm, n = 791, *N* = 4; *P* = 0.95; *Glialcam*-null KA: 1.55 ± 0.02 µm, n = 1002, *N* = 5 vs *Glialcam*-null KA recovery: 1.50 ± 0.02 µm, n = 909, *N* = 4; *P* = 0.99, nested one-way ANOVA; **Fig. 4**). To conclude, we found that astrocyte processes in 3-5-month-old *Glialcam*-null mice are comparable with regard to process thickness to age-matched wild-type mice. In addition, we found that seizure activity does not lead to acute swelling of astrocyte processes in wild-type or in *Glialcam*-null mice.

## DISCUSSION

We investigated whether two pathological features of MLC, vacuolization of cerebellar white matter and thickness of astrocyte cell processes abutting blood vessels, aggravate upon severe seizure activity. Additionally, we tested to what extent myelin vacuoles develop in wild-type mice undergoing severe seizure activity. Seizure activity in mice was induced by injecting the chemoconvulsant KA. Since wild-type and MLC mice respond differently to a single dose of KA (Dubey et al., 2018), we used a regime of repeated low dose KA injections until a mouse developed severe seizures (Tse et al., 2014). We demonstrated the efficacy of this approach in generating severe seizure activity in all mice, regardless of genotype. Cumulative seizure severity over the total duration of the experiment was comparable between wild-type and *Glialcam*-null mice, meaning that seizure intensity can be considered equal. Surprisingly, the induced severe seizure activity did not affect myelin vacuolization or astrocyte process thickness in either *Glialcam*-null or wild-type mice, implying that a single, short period of intense seizure activity does not acutely worsen pathological features of MLC.

In our study we did not observe a difference in seizure behavior between *Glialcam*-null mice and wild-type mice. In 8-to 12-month-old *Glialcam*-null mice, a single injection of 10mg / kg KA leads to the development of severe seizures (Dubey, Brouwers et al, 2018), while a similar single injection only leads to modest seizures in wild-type mice. The fact that no difference in seizure behavior was observed in the present study can have several explanations. One possibility is that *Glialcam*-null mice are specifically more sensitive to acute intense stimulation (a single high dose of KA) than wild-type mice, but that the sensitivity to repeated low dose injections is similar. Alternatively, our current behavioral scoring (time to first Racine score 4/5 seizure and maximum seizure score per 5 min interval) might not be fine-grained enough to pick up a subtle difference in seizure threshold or behavioral response. Finally, the absence of a difference in seizure behavior could be due to the young age of the mice. In our previous study, we used 8-to 12-month-old mice (Dubey et al., 2018), while in the current study we used slightly younger mice (3-to 5-month-old). We reasoned that the relatively lower number of vacuoles present in the younger age group would be beneficial for measuring additive seizure induced vacuolization. However, the lowered seizure threshold in MLC mice might only develop at an older age. Nonetheless, it should be noted that stimulation-induced potassium dynamics in acute brain slices from MLC mice was already disturbed in slices from 3-to 5-month-old animals (Dubey et al., 2018). Therefore, if disturbed potassium dynamics underlies the decreased seizure threshold we would expect the seizure threshold to be lowered already in 3-to 5-month-old mice. A definite confirmation of this hypothesis could be achieved by injecting a single high dose of KA in 3-to 5-month-old mice.

Histologically, MLC is characterized by myelin vacuoles and swollen astrocyte endfeet. How or why these features arise is incompletely understood. We have hypothesized that the impairment of astrocyte volume regulation primarily leads to astrocyte swelling. This would subsequently hamper the efflux pathway of excess potassium and water from myelinated axons, thereby leading to swelling and vacuolization of myelin (Min & van der Knaap, 2018; van der Knaap, Boor, & Estevez, 2012). Since both astrocyte swelling and myelin vacuolization are thought to be related to neuronal activity (Blanz et al., 2007; MacVicar, Feighan, Brown, & Ransom, 2002; Menichella et al., 2006), we hypothesized that these features would be aggravated by seizure activity. However, our results find no indication for the occurrence of these morphological changes following a period of excessive seizure activity. This surprising lack of effect might be due to the duration of seizure activity or to the time between seizure activity and brain fixation. In the following section we will discuss first myelin vacuolization and then astrocyte end-foot thickness in more detail.

The expected timeline after seizures at which myelin vacuolization might appear and resolve is unclear. In our study we examined vacuolization directly following 1 hour of severe seizure activity, but were unable to observe increased vacuolization. Early experiments on dorsal root stimulation of the frog suggest acute development of myelin vacuoles on a similar timescale. Wurtz & Ellismann directly stimulated frog dorsal root fibers for 15 minutes at a frequency of 20-50 Hz. Tissue fixation immediately following stimulus termination revealed prominent vacuolization of myelin. Importantly, a 30-minute stimulation-free period was sufficient to recover from these structural changes in myelin. We did not observe similar acute development of myelin vacuoles in the brain. It is important to mention that tissue fixation in our experiments did not always immediately follow a severe seizure. Therefore it is possible that we missed the peak of vacuolization, if vacuolization indeed develops and resolves acutely. Equally possible is that the local stimulation of the frog fiber had a greater impact on the myelin than the globally induced neuronal activity in our study. Although our findings suggest that there is no acute vacuolization of myelin in the MLC or healthy brain after a period of severe seizures, our results leave open the possibility that vacuoles will develop over time following seizure activity. This would be in line with observations in the Cx32/Cx47 double knockout (dKO) mouse (Menichella et al., 2006). Activity-dependent optic nerve vacuolization in the Cx32/Cx47 dKO mouse was observed 48 hours after stimulation with a cholera toxin injection, suggesting longer term effects of excessive neuronal activity on myelin integrity. In line with this, we observed a small increase in myelin vacuolization in *Glialcam*-null mice that were allowed to recover from seizures for 6 hours following the repeated low-dose KA injections. Therefore, it might be interesting to include later time points in a feature study on seizure induced myelin vacuolization.

The cerebellum is susceptible to the development of vacuoles. This is obvious from our earlier studies in mouse models for MLC (Bugiani et al., 2017; Dubey et al., 2015), but also in the spontaneous epileptic rat (SER) where myelin vacuoles mainly develop in the cerebellum and brainstem (Inui et al., 1990). However, while brain edema and histological alterations following KA-induced seizures have been described in several brain regions (Sperk et al., 1983), whether such dramatic alterations occur in the cerebellar white matter is unclear from previous studies. In this respect it is important to note that visual inspection of the corpus callosum, a structure characterized by mild vacuolization in MLC mice, also did not reveal clear KA-induced vacuolization under any of the conditions in this study (data not shown). Another structure clearly involved in KA-induced seizures is the hippocampus. However, since vacuolization of hippocampal structures in the context of MLC has not been described, we did not study hippocampal morphology following KA-induced seizures in the present study.

We investigated whether thickness of astrocyte processes abutting blood vessels is increased upon severe seizure activity. As with myelin vacuolization, we found no activity-dependent thickening of astrocyte processes following 1 hour of KA induced severe seizure activity. Astrocyte swelling in response to KA-induced seizure activity has been reported to occur in rats. A single injection of 10mg / kg KA led to marked perineuronal and perivascular astrocyte swelling within 1 hour after the onset of severe seizure activity and was most pronounced after 2 hours (Lassmann et al., 1984). Therefore, astrocyte cell swelling can take place on short time scales. Our observations do not confirm this. We cannot rule out that total severity of seizure activity has been milder in our experiments compared to the study by Lassmann and colleagues, since a detailed behavioral scoring of seizure severity is lacking in the Lassmann study. Additionally, our analysis focused specifically on the thickness of perivascular astrocyte processes, and did not study overall astrocyte cell swelling. The reason for focusing on perivascular process thickness was a clear increase in this parameter in MLC mice observed in earlier studies (Bugiani et al., 2017; Dubey et al., 2015). Process thickness was assessed using a GFAP staining of the tissue. The intermediate filament GFAP is part of the astrocyte cytoskeleton (Middeldorp & Hol, 2011). While thickness of GFAP processes might reflect the chronic swelling of astrocyte processes in MLC, it might not be the best measure for acute swelling of astrocyte processes. The filling of the swollen astrocyte process with GFAP positive cytoskeleton might take time. Therefore, future studies would benefit from cytosolic expression of fluorophores in the astrocyte, followed by morphometric analysis of the whole astrocyte, to more specifically address whether acute astrocyte swelling occurs (Suzuki et al., 2003) (Rosic et al., 2019). For example, using this method it was found that KA-induced severe seizure activity leads to acute development of astrocyte vacuoles, peaks upon 4 hours after seizure termination and persists for up to 3 days (Guo, Zou, & Wong, 2017).

Perivascular astrocyte processes of *Glialcam*-null mice were previously reported to be significantly thicker compared to wild-type mice (Bugiani et al., 2017). We did not observe this difference in our study. This may be due to the young age of the wild-type mice, which at 3 months were notably younger than the 9-to 12-month-old mice used by Bugiani et al. (2017). Age-dependent changes in astrocytes are not uncommon. In addition to morphological development, astrocytes undergo changes in volume regulation (Rodriguez-Arellano, Parpura, Zorec, & Verkhratsky, 2016) (Kolenicova et al., 2020). Mouse astrocytes swell upon exposure to pathological concentrations of potassium but the extent with which they swell fluctuates between 3 and 18 months. The percentage of swelling even seems to halve between 3 and 9 months (Kolenicova et al., 2020). It may therefore well be possible that astrocyte endfeet thickness also fluctuates with age.

One important open question regarding the link between myelin vacuoles and seizures is what the functional impact of vacuolization is on neuronal signal transmission. It is presently unclear whether vacuolization alters action potential propagation and, if so, what the direction of this effect is. Temporary hyperexcitability of the axon might result from potassium accumulation in the periaxonal space underneath the myelin because of an obstructed potassium exit pathway, leading to axonal depolarization (Brazhe et al., 2011). If prolonged, this might lead to conduction failures or even complete conduction block (Brazhe et al., 2011). Early functional experiments in frog nerve show that swelling-induced alteration of myelin integrity leads to slowing of conduction velocity, as assessed by measurement of the compound action potential (Wurtz & Ellisman, 1986). However, to our knowledge, experimental studies directly linking the presence of myelin vacuoles to axonal action potential properties in mammalian axons are lacking. To fully understand the impact of vacuolization on neuronal signal transmission such studies are required. Mathematical modelling of periaxonal potassium dynamics and myelin properties have already yielded interesting results (Brazhe et al., 2011).

In conclusion, in our study we show that repeatedly injecting a low dose of KA provides the opportunity to regulate seizure development and to generate an equal amount of severe seizure activity in both wild-type and epilepsy-sensitive *Glialcam*-null mice. Histochemical characterization of these mice following seizures shows that KA-induced severe seizure activity has no acute effect on the development of myelin vacuolization in *Glialcam*-null or wild-type mice. We did observe a small increase in myelin vacuoles in *Glialcam*-null mice several hours after seizures had stopped, which was absent in wild-type mice. This might indicate that activity dependent formation of myelin vacuoles takes place over a longer timescale than investigated here. Finally, we find that astrocytes do not undergo seizure induced morphological changes in the form of thickened processes abutting blood vessels. Thus, our study suggests that the two major pathological features of MLC, myelin vacuolization and swollen astrocyte endfeet, do not aggravate acutely in response to severe seizure activity.

